# Outer membrane vesicles derived from *Salmonella* Typhimurium can deliver *Shigella flexneri* 2a O-polysaccharide antigen to prevent lethal pulmonary infection in mice

**DOI:** 10.1101/2021.02.01.429291

**Authors:** Huizhen Tian, Biaoxian Li, Yuxuan Chen, Kaiwen Jie, Tian Xu, Zifan Song, Xiaotian Huang, Qiong Liu

**Author notes:** **Corresponding authors: Qiong Liu Email address**.

## Abstract

The threat to health from shigellosis has become quite serious in many developing countries, causing severe diarrhea. *Shigella flexneri* 2a (*S. flexneri* 2a) is the principal species responsible for this endemic disease. Although there have been multiple attempts to design a vaccine against Shigellosis, one that is effective has not yet been developed. Lipopolysaccharide (LPS) is both an essential virulence factor and a protective antigen against *Shigella*, due to its outer domain, termed O-polysaccharide antigen. In the present study, *S. flexneri* 2a O-polysaccharide antigen was innovatively bio-synthesized in *Salmonella* and attached to core-lipid A by the ligase WaaL, and thus purified outer membrane vesicles (OMVs) were used as a vaccine for subsequent research. Here, we identified the expression of the heterologous polysaccharide antigen and described the isolation, characterization, and immune protection efficiency of the OMV vaccine. The expression of *S. flexneri* 2a did not affect the formation of *Salmonella* OMVs or their cytotoxicity. Furthermore, the results of the animal experiments indicated that immunization of the mice both intranasally and intraperitoneally with the OMV vaccine induced significant specific anti-Shigella LPS antibodies in both vaginal secretions and fluid from bronchopulmonary lavage, in addition to within sera. The OMV vaccine immunized by both routes of administration provided significant protection against virulent *S. flexneri* 2a infection, as judged by a serum bactericidal assay (SBA), opsonization assay, challenge experiment, and pathological analysis. The present study firstly indicates that the proposed vaccination strategy represents a novel and improved approach to control Shigellosis by the combination of bioconjugation of *Salmonella* glycosyl carrier lipid and OMV. In addition, the strategy of genetic manipulation described here is ideally suited for designs based on other *Shigella* serotypes, allowing the development of a *Shigella* vaccine with broad-protection.

## Introduction

Shigellosis continues to be a leading cause of severe inflammatory diarrhea in many developing countries, thought to cause approximately 165 million cases each year, predominantly in children under 5 years of age ^1,2^. Among all serotypes, *Shigella flexneri* (*S. flexneri*) is the major cause of bloody diarrheal disease in humans and is also an important pathogen in higher primates in a numer of settings ^3,4^. As occurs with other enteric Gram-negative bacteria, *Shigella* can invade intestinal cells, the infection of which should result in a pro-inflammatory response ^5^. O-antigen (O-Ag) chains, a component of lipopolysaccharide (LPS) molecules, contribute to *Shigella* virulence and infection ^6^. O-antigen chains are formed by oligosaccharide repeating units (RUs) ^3,7^ that bear a linear tetrasaccharide backbone consisting of three L-rhamnose residues and an N-acetyl-D-glucosamine residue ^8^ which are encoded by O-antigen gene clusters approximately 16,000 bp in length and located between the *galF* and *gnd* genes ^9^ (**Fig. 1A**).

**Figure 1.**
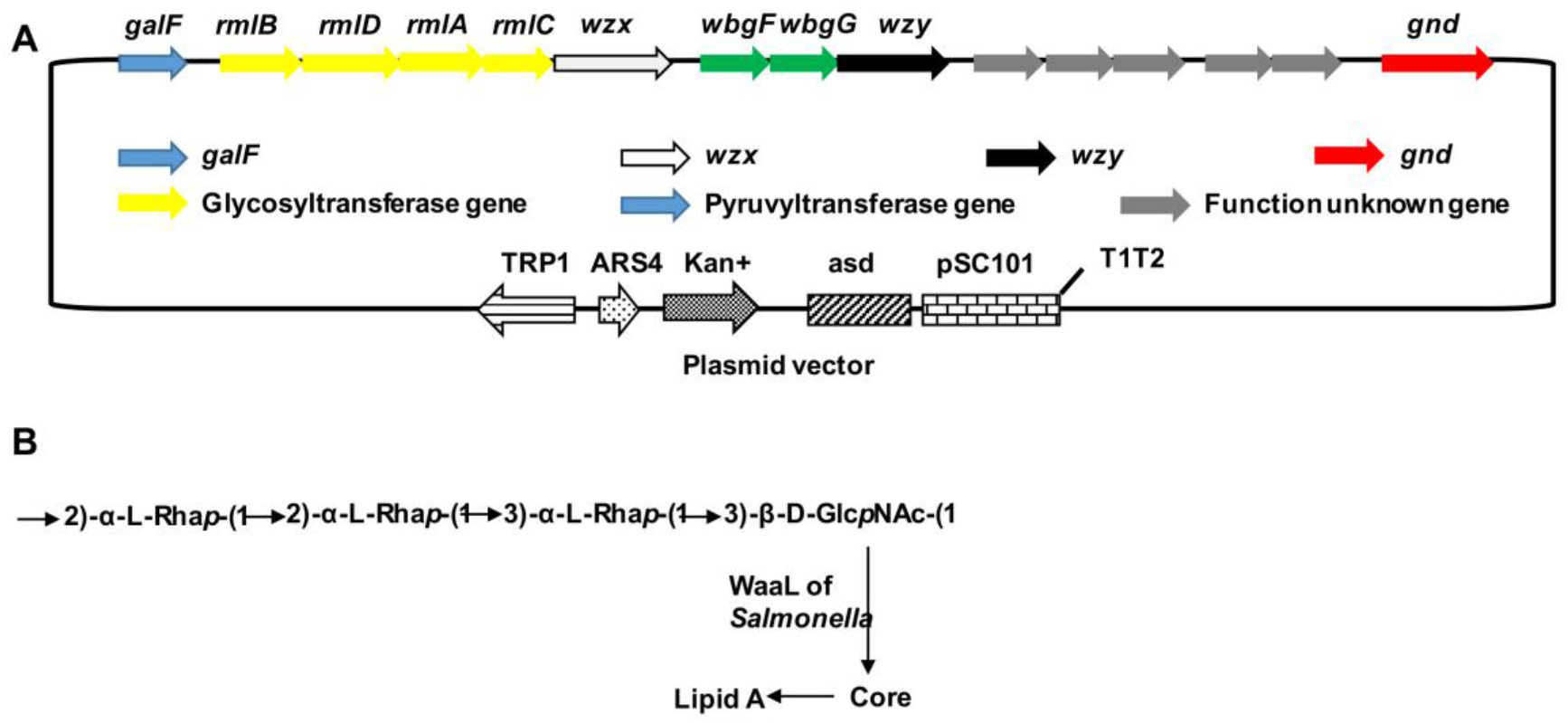
(A) Construction of plasmid expressing *S. flexneri* 2a O-antigen polysaccharide. The origin for replication was pSC101 as the replicon. The complete O-antigen cluster was cloned in this plasmid then expressed. (B) Schematic molecular model of the structure of *S. flexneri* 2a O-antigen and the principle of expression in *S.* Typhimurium.

Vaccination is a pivotal aspect of the strategy for controlling shigellosis ^10^. The humoral immune response, including systemic and mucosal responses, is a major component of protective immunity against *Shigella* infection ^11–13^ and available data suggests that the presence of type-specific serum antibodies recognizing the O-antigen of the LPS of *Shigella* is associated with protection against shigellosis ^12^, even producing some cross-reactivity ^14^. However, serum antibodies alone are not sufficient to predict whether someone has protection against shigellosis ^15^. Mucosal immunity plays a critical role in the mechanism of protection against *Shigella* infections, as established in previous studies ^16,17^. Levels of antibody-secreting cells (ASCs), especially those that secrete immunoglobulin A (IgA) antibodies, could be consistent with the degree of protection capacity for a potential vaccine ^18^. A mouse model has recently been established in studies that evaluated the safety and efficacy of *Shigella* vaccines ^19,20^. Minimal reactogenicity and significant protection efficacy were observed in vaccinated mice, and were found to be safe and able to protect volunteers and primates ^21,22^.

Outer membrane vesicles (OMVs) produced by Gram-negative bacteria contain biologically active components such as proteins and LPS, which perform diverse biological processes including participation in the secretory pathway, bacterial infection, and OMV-based vaccines ^23–25^. Parenteral vaccines are often ineffective in stimulating the mucosal immune response, instead most effectively elicited by antigens at mucosal surfaces ^26,27^. It has been established that OMVs as nano-particle vaccines induce mucosal immunity and protection against *Shigella* and other intestinal bacteria ^28,29^. A previous study has demonstrated that immunization against *S.* Typhimurium-derived OMVs stimulates a mucosal immune response and confers protection against *Salmonella* infection ^30^. OMVs derived from intracellular bacteria, especially *S.* Typhimurium, represent a platform for the delivery of heterologous antigens. However, the majority of antigens are immunogenic proteins ^31^. Recently, two articles have confirmed that heterologous polysaccharides acting as important protection antigens could be incorporated into OMVs and induce immune activation, with glycoengineered OMVs demonstrating considerable potential as a vaccine due to good efficacy and safety of the OMV platform ^32,33^.

In the present study, the feasibility of delivering a vector based on OMVs from *S.* Typhimurium carrying *S. flexneri* 2a O-polysaccharide antigen was explored and a potential vaccine against *Shigella* infection evaluated (**Figs. 1A and 1B**). Efficient delivery vehicles were fabricated by judging the correct balance of immune response between homologous and heterologous antigens ^34,35^. Therefore, non-essential antigens which displayed a strong immune response to the host and disrupted the recognition of delivery antigens by antigen-presenting cells (APCs) were omitted, such as flagellin proteins FliC and FljB and major porins including OmpA, OmpC, and OmpD. To ensure the display of heterologous O-antigens, the *rfbP* gene was knocked out which enabled the expression of *S. flexneri* O-antigen on core-lipid A. The immune response and protection afforded by this *Shigella* glycoconjugate-OMV vaccine were evaluated in a mouse model, establishing a novel and improved approach for the prevention of Shigellosis.

## Materials and Methods

### Construction of plasmids and mutants

All strains and plasmids utilized in the present study are listed in Table 1. *S.* Typhimurium, *Escherichia coli* (*E. coli*), and *S. flenxeri* 2a were cultured in Luria-Bertani (LB) broth or agar (Difco, Detroit, MI) at 37°C. Diaminopimelic acid (DAP) (50 μg/ml) was added for the growth of Δ*asd* strains ^36^. LB agar containing 5% sucrose was used for *sacB* gene-based counterselection for the generation of mutant strains.

**Table 1.**
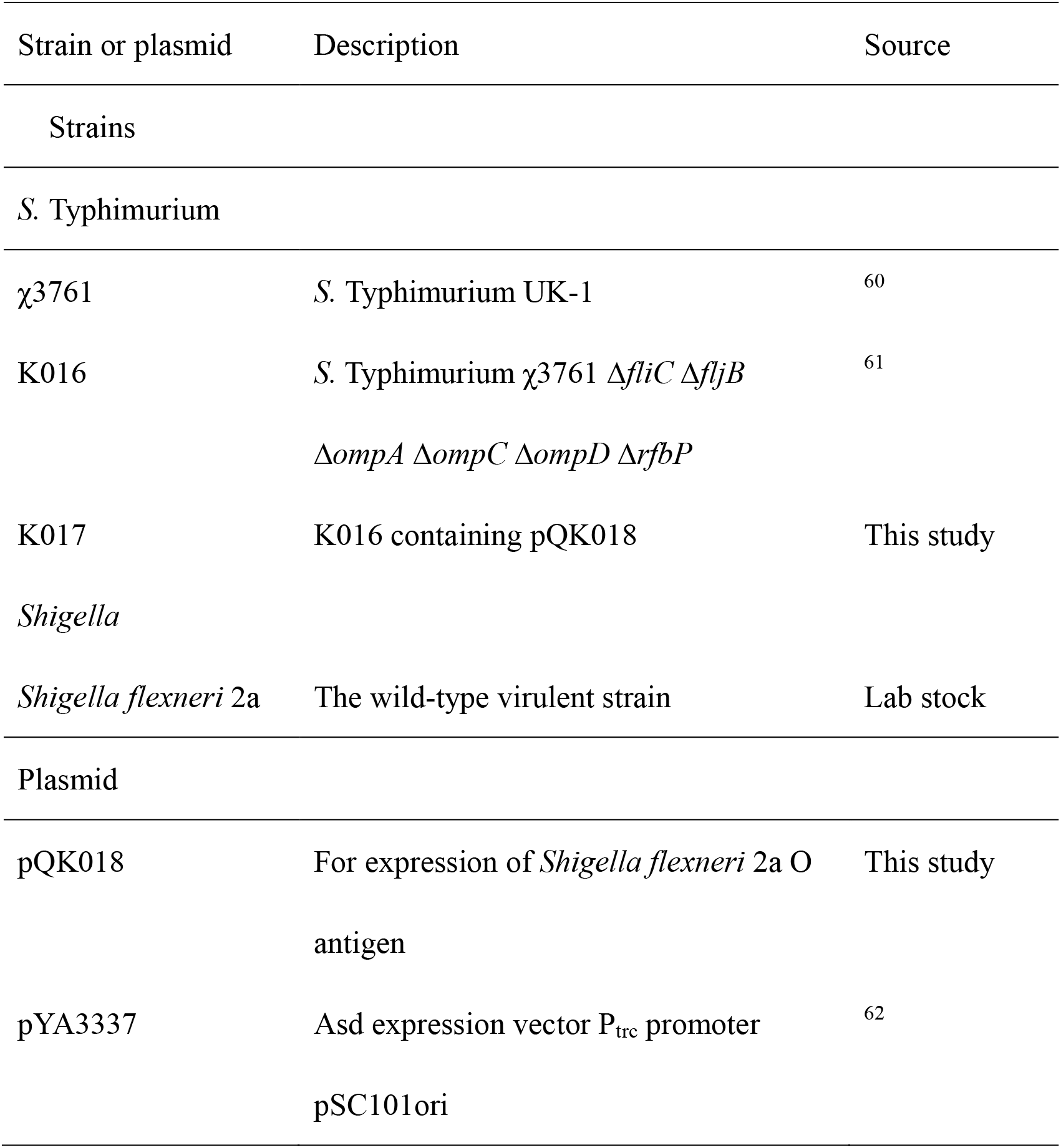
Bacterial strains and plasmids used in this study.

| Strain or plasmid | Description | Source |
| --- | --- | --- |
| Strains |  |  |
| *S. Typhimurium* |  |  |
| χ3761 | *S. Typhimurium* UK-1 | 60 |
| K016 | *S. Typhimurium* χ3761 $\Delta fliC$ $\Delta fljB$<br>$\Delta ompA$ $\Delta ompC$ $\Delta ompD$ $\Delta rfbP$ | 61 |
| K017 | K016 containing pQK018 | This study |
| *Shigella* |  |  |
| *Shigella flexneri* 2a | The wild-type virulent strain | Lab stock |
| Plasmid |  |  |
| pQK018 | For expression of *Shigella flexneri* 2a O antigen | This study |
| pYA3337 | Asd expression vector P<sub>trc</sub> promoter<br>pSC101ori | 62 |

The primers used in the present study are listed in Table 2. DNA manipulation was conducted, as described elsewhere ^37^. Transformation of *E. coli* and *S.* Typhimurium was performed by electroporation. Transformants were selected on LB agar plates containing appropriate antibiotics.

**Table 2.**
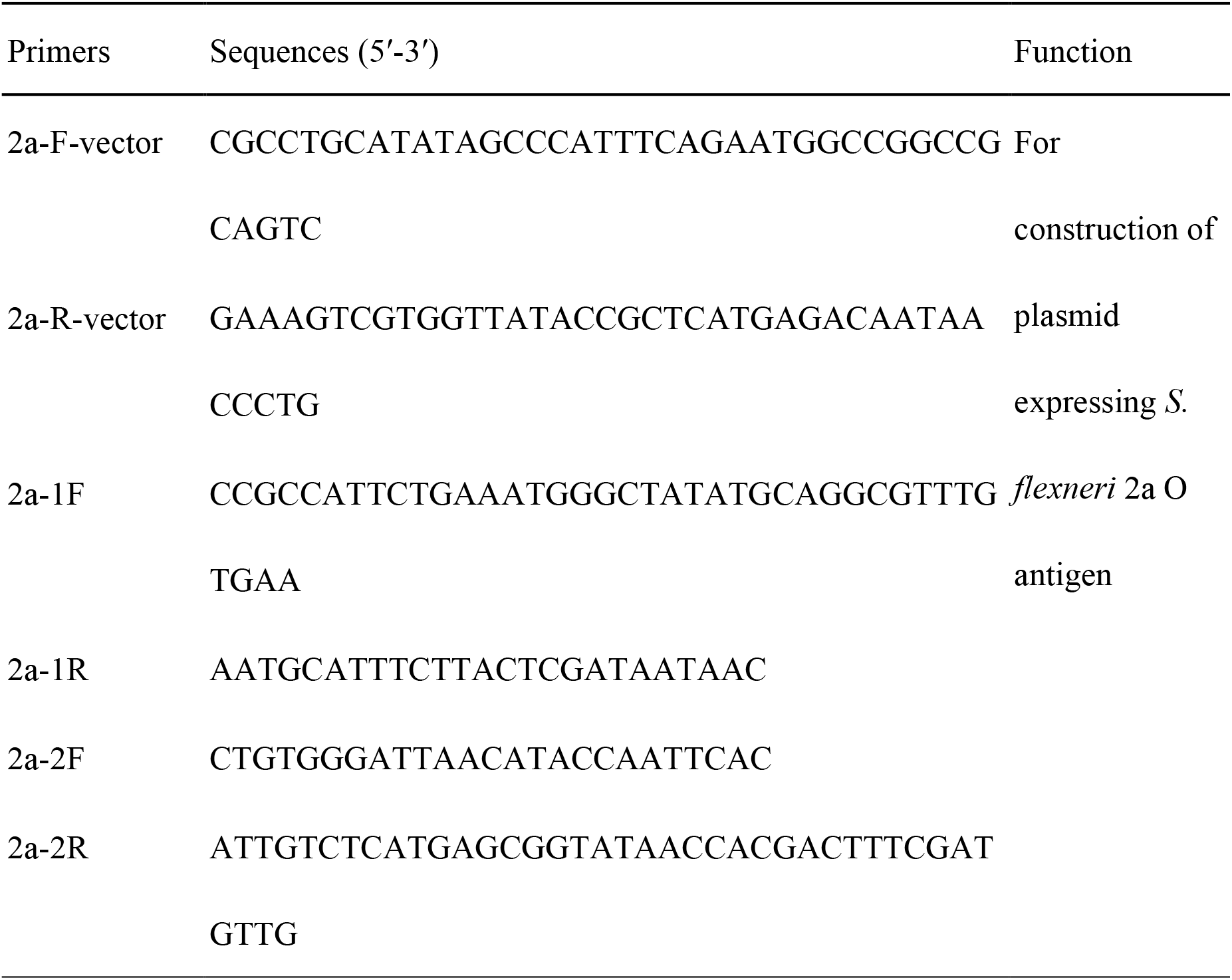
Primers used in this study.

For construction of the expression plasmid pQK018, *S. flexneri* 2a O-antigen gene clusters were divided into two parts then amplified by two pairs of primers, 2a-1F/2a-1R and 2a-2F/2a-2R, using the *S. flexneri* 2a genome as a template. An approximate 6000-bp fragment of the plasmid vector was amplified by primers, 2a-F-vector/2a-R-vector. Those fragments overlapped and were assembled using a Gibson assembly kit (New England Biolabs, Beverly, MA) in accordance with the manufacturer’s protocol, resulting in expression of pQK018 (**Fig. 1A**). Furthermore, the *S. flexneri* 2a O-antigen expression plasmid pQK018 was transferred into the previously constructed mutation K016, for the successful construction of a mutant strain capable of expressing *S. flexneri* 2a O-antigen (K017) for subsequent purification of the OMV vaccine.

### OMV purification and measurement of concentration

OMVs were isolated from *Salmonella* essentially as described in a previously published protocol ^31^. Total protein concentration in the OMVs was measured using a bicinchoninic acid (BCA) protein assay (Thermo Pierce, Rockford, IL), in accordance with the manufacturer’s protocol.

### Cryo-electron microscopy, lipopolysaccharide assays, and Western blot analysis

Cryo-electron microscopy was performed as described previously ^38^. Lipopolysaccharide profiles of the OMVs or strains were examined in the same manner as the method used for *Salmonella* ^39^. After separation of LPS in the samples by SDS-PAGE, the different bands were blotted to nitrocellulose membranes. The membranes were incubated with blocking solution (5% skim milk in Tris-buffered saline) for 2 h, then incubated with rabbit polyclonal antibodies specific for *S. flexneri* 2a O-antigen (BD company, Franklin Lakes, NJ) at room temperature. An alkaline phosphatase-conjugated goat anti-rabbit immunoglobulin G secondary antibody (Sigma-Aldrich, St Louis, MO) at 1:10,000 dilution was added and incubated at room temperature for 1 h. Immunoreactive bands were detected by the addition of BCIP (5-bromo-4-chloro-3-indolylphosphate)-nitroblue tetrazolium solution (Sigma-Aldrich). The reaction was stopped after 2 min by washing the blots with large volumes of deionized water.

### Cytotoxicity of OMVs toward macrophages

RAW264.7 murine macrophages were obtained from the Cell Bank of the Chinese Academy of Sciences (Shanghai, China). The cells were maintained at 37°C in Dulbecco’s modified Eagle medium (DMEM, Gibco BRL, Gaithersburg, MD) supplemented with 10% fetal bovine serum (FBS, Hyclone, Logan, UT) in an atmosphere containing 5% CO_2_. The specific cytotoxicity of the OMV vaccine or OMV vector in macrophages was conducted by seeding RAW264.7 cells in 24-well plates (5 × 10^5^ cells/well) then exposing them to different concentrations of OMVs, ranging from 3.075 μg/ml to 100 μg/ml. The plates were incubated at 37°C in 5% CO_2_ for 24h. The supernatants of each well were collected then assessed using a Multitox-Fluor Multiplex Cytotoxicity Assay (Promega, Madison, WI), in accordance with the manufacturer’s instructions. Supernatants of cells cultured without OMVs or treated with cell lysis solution (Biovision, California, USA) were used as negative and positive controls, respectively. The value for each experiment was determined from the mean of wells measured in triplicate. In addition, three independent experiments were performed.

### OMV immunization protocol in mice

Research in animals was conducted in compliance with the Animal Welfare Act and regulations related to experiments involving animals (Ya’an, China; Approval No. 2011-028). Principles stated in the Guide for the Care and Use of Laboratory Animals were followed. All experimental protocols were approved by Sichuan Agricultural University. All efforts were made to minimize any suffering of animals during the experiments.

Six-week-old female BALB/c mice (16-22 g) were purchased from Dashuo Biotechnology Co., Ltd. (Chengdu, China), and used in immunization trials, in which two immunization methods were used. Suitable groups of 10 mice each were immunized on days 0 and 30 with 20 μg OMVs in 10 μl PBS per mouse by intranasal administration or 5 μg OMVs in 100 μl PBS buffer per mouse by intraperitoneal injection. Ten μl of PBS administered nasally or 100 μl PBS by intraperitoneal injection served as negative controls in the two methods of immunization, respectively. Blood samples were collected by orbital sinus puncture and vaginal secretions collected by repeated flushing and aspiration of a total of 0.15 ml of the same buffer at 4 and 8 weeks after the first immunization. Lung lavage samples were collected 4 weeks after the booster immunization in the groups with intranasal administration. Following centrifugation, serum was obtained, all samples were preserved at −80°C.

### Challenge Experiment

To determine the cross-protective efficacy of the recombinant OMV vaccine against wild-type parental or heterologous *Salmonella*, mice were infected with approximately 10^9^ colony-forming units (CFU) (10,000-fold LD_50_) of *S.* Typhimurium, 10^9^ CFU (10,000-fold LD_50_) of *S.* Choleraesuis and 10^9^ CFU (10,000-fold LD_50_) of *S.* Enteritidis in 20 μl PBS with 0.01% gelatin (BSG buffer), by oral administration 5 weeks after booster immunization, respectively. Challenged mice were monitored for three weeks and survival and death were recorded daily. During this period, infected mice were closely monitored twice per day for their clinical appearance. If obvious suffering was observed (weight loss, ruffled fur, hunched posture, decrease in appetite, weakness/inability to obtain feed or drink water, lethargy, or morbidity), the mice were immediately euthanized using CO_2_ in a custom flow metered chamber. At the conclusion of the animal experiments, all remaining mice were euthanized with CO_2_. The animal experiments were performed twice. Data were combined prior to analysis.

To evaluate the protection efficacy of the vaccine against virulent *Shigella* 5 weeks after the booster immunization, a bacterial suspension (20 μl) was administered drop-wise on the external nares of each mouse using a 100 ml Hamilton syringe. The mice were challenged with a lethal dose (approximately 10^6^ CFU, 100-fold LD_50_) of *S. flexneri* 2a. Challenged mice were monitored daily for 30 days. Animal care was consistent with the previous procedures. This animal experiment was performed twice, and data combined prior to analysis.

### Splenic lymphocyte proliferation

Mice were euthanized 7 days after primary immunization (3 mice per group) and their spleens aseptically removed. Spleen cell suspensions were obtained then passed through a 70 μm sterile cell strainer (Fisher Brand, Houston, TX; 1 spleen/strainer). Erythrocytes were lysed with RBC lysis buffer (eBioscience, San Diego, CA). Splenocytes were resuspended in RPMI 1640 culture medium supplemented with 5% FBS. The suspended cells were seeded in 96-well culture plates (3 × 10^5^ cells/well) then incubated in a 5% CO_2_ incubator at 37°C. For the experimental test groups, cells were cultured with *S. flexneri* 2a LPS or *S.* Typhimurium LPS (30 ng/well). As a positive control, cells were cultured with phytohemagglutinin (PHA, 10 μg/ml), and with sterile culture medium as a negative control. After 72 h of incubation, 20 μl of MTT (5 mg/ml in PBS) were added to each well. After 4 h of incubation at 37°C in a 5% CO_2_ incubator, supernatants were collected after centrifugation at 200 g for 5 min. Formazan crystals were dissolved by the addition of 100 μL DMSO per well and agitation at 37°C for 10 min. The OD at 490 nm was measured using a microplate reader. Cell proliferation was calculated as a stimulation index (SI), defined as the ratio of the mean absorbance of triplicate wells of cells stimulated with PHA or LPS to that the negative control.

### Measurement of cytokine production by stomach tissue and mouse splenocytes

Mouse splenocytes were isolated to evaluate the Th1/Th2 polarizing effects of immunization through measurement of the levels of IFN-γ, IL-12 (p40), IL-4, IL-13, IL-6, and TNF-α. The splenocytes of nine mice that had received booster immunization with the corresponding antigens were obtained and stimulated for 24 hours with 6 μg/ml LPS isolated from *S. flexneri*, as previously reported ^40^. Supernatants from stimulated cells were collected and cytokines produced by the stimulated cells measured by ELISA. The detailed ELISA protocol is described below. Briefly, 96-well plates were coated with monoclonal anti-IFN-γ, anti-IL-12 (p40), anti-IL-4, anti-IL-13, anti-IL-6, or anti-TNF-α antibodies (BD Biosciences, Mountain View, California, USA). After blocking with 1% BSA in PBS, the samples were added to duplicate wells and incubated overnight at 4°C. The wells were washed and incubated with biotinylated monoclonal anti-INF-γ, anti-IL-12 (p40), anti-IL-4, anti-IL-13, anti-IL-6, or anti-TNF-α antibodies (BD Biosciences, Billerica, Massachusetts, USA). Horseradish peroxidase (HRP)-labeled anti-biotin antibody (Vector Laboratories, Burlingame, California, USA) was then added to each well. The reaction was developed by the addition of 3,3′,5,5′-tetramethyl-benzidine (Moss Inc., Pasadena, California, USA) and stopped with 0.5 N HCl. Standard curves were generated using mouse recombinant (r) IFN-γ, IL-12 (p40), IL-4, IL-13, IL-6, and TNF-α.

### Opsonization assay

An opsonization assay was performed using a method described in a previous study ^41^. Briefly, mouse peritoneal macrophages were harvested by flushing the peritoneal cavity of BALB/c mice with precooled PBS. Macrophages collected in the fluid were centrifuged and resuspended in prewarmed RPMI (Gibco BRL, Gaithersburg, MD) supplemented with 10% FBS. Approximately 5 × 10^5^ cells were added to the wells of 12-well plates and cultivated at 37°C in an atmosphere containing 5 % CO_2_ for 2h. Non-adherent cells were removed and the medium replaced with fresh medium, then incubated at 37°C overnight. Prior to inoculation, a log-phase culture of 10^6^ *S. flexnari* 2a in PBS was incubated with immunized or non-immunized sera (negative control) for 1.5h at 37°C. Macrophages were infected with 5 × 10^6^ opsonized bacteria (MOI of 10 bacteria/macrophage) and incubated for 30 min. Macrophages were washed with PBS then incubated with gentamicin (50 μg/ml) in RPMI for a further 30 min. The macrophages were washed then incubated with RPMI for 0 or 60 min, after which they were lysed with 0.1% Triton X-100. The number of *S. flexneri* were determined by CFU count on LB plates.

### Serum bactericidal assay (SBA)

A serum bactericidal assay (SBA) was performed using the protocol described previously, with some modifications ^42^. Briefly, *S. flexneri* 2a were cultured in LB medium to log phase (OD=0.2), diluted 1:10,000 in SBA buffer (PBS + 0.5% BSA) to approximately 10^3^ CFU/ml, then distributed into sterile polystyrene U-bottomed 96-well plates. In each well, a total volume of 50 μl, containing 12.5 μl of the bacterial cells, 25 μl of 2-fold increasing concentrations of mice serum (starting at a 1:100 dilution), and 12.5 μl of baby rabbit complement (Sigma-Aldrich, St Louis, MO). The sera were heated to 56°C for 30 min to inactivate any endogenous complement. To evaluate the possible nonspecific inhibitory effects of rabbit complement or mouse serum, the bacteria were also incubated with either the same tested sera plus heat-inactivated complement or SBA buffer and activated complement. The plates were incubated at 37°C for 90 min, and 10 μl of the sample from each well spotted onto LB agar plates, the resultant CFUs counted on the following day. Bactericidal activity was calculated from the proportion of CFUs counted in each dilution of serum with active or inactive complement compared with CFUs of the same serum dilutions with no complement. Each sample and control were tested in triplicate.

### Quantitative ELISA

The methods of purification of LPS from *S. flexneri* 2a were described in a previous study ^27^. The antibody response was analyzed by quantitative ELISA. One μg of LPS suspended in 100 μl sodium carbonatebicarboante coating buffer (pH 9.6) was used to coat 96-well plates and then incubated overnight at 4°C. Standard curves of each isotype were created. The concentration of antibody was quantified in plates coated in triplicate with two-fold dilutions of an appropriate purified mouse Ig isotype standard (IgG and IgA; BD Biosciences), starting at 0.5 μg/μl. Each plate was washed 3 times with PBS containing 0.1% Tween 20 (PBST) then blocked with 2% BSA solution for 2 h at room temperature. A 100 μl volume of suitably diluted sample was added to individual wells in triplicate and incubated for 1h at room temperature. After washing with PBST, biotinylated goat anti-mouse IgG and IgA (Southern Biotechnology Inc., Birmingham, AL) were added to each well. The wells were then developed with a streptavidin-alkaline phosphatase conjugate (Southern Biotechnology Inc.) then measured using a p-nitrophenylphosphate substrate (Sigma-Aldrich) in diethanolamine buffer (pH 9.8). The depth of color (absorbance) was measured at 405 nm using an automated ELISA plate reader (model EL311SX; Biotek, Winooski, VT). The final Ig isotype concentration in the antibody sample was calculated using an appropriate standard curve. A log-log regression curve was calculated from at least four dilutions of the isotype standard.

### Histology and Pathological scores

Animals were euthanized by inhalation of CO_2_. The lungs were removed then perfused with 10% buffered formalin phosphate (Fisher, Pittsburgh, PA), dehydrated, then processed in paraffin. Sections were cut at 3 mm and intervals then stained with hematoxylin and eosin, or Giemsa.

The lung injury score was recorded by a pathologist blinded to the grouping on a 0 to 4 point scale, based on the following parameters: infiltration of inflammatory cells, epithelial shedding, loss of barrier integrity, and goblet cell hyperplasia ^43^.

### Statistical analysis

All experiments were conducted in triplicate. A one-way analysis of variance (ANOVA) was performed to determine the statistical significance of differences between mean values of various experimental and control groups. Data were expressed as means ± standard deviation. Means were compared using a least significant difference test. *P* < 0.05 was considered significantly different. All data were analyzed with GraphPad Prism version 5.01.

## Results

### Identification of *Salmonella* OMV vaccine delivering *Shigella* O-polysaccharide antigen

To construct a plasmid expressing the complete *S. flexneri* 2a LPS O-antigen, *S. flexneri* 2a O-antigen gene cluster, located between the *galF* and *gnd* genes and approximately 16,000 bp in size (**Fig. 1A**), was cloned by PCR. The low copy plasmid was selected as the vector plasmid pYA3337, whose origin of replication was pSC101 ^44^. In the present study, pQK018 expressing the *S. flexneri* 2a O-antigen was constructed using a Gibson assembly kit (New England Biolabs, Beverly, MA) (**Fig. 1B**). The identity of the expression plasmid was in found in the TOP10 *E. coli* strain (Invitrogen, Carlsbad, CA) due to its rough phenotype of LPS. The *S. flexneri* 2a O-antigen could be expressed in the TOP10 strain, as it was visible in SDS-PAGE gels using silver staining (Supplemental **Fig. 1**).

Purified *Salmonella*-derived OMVs delivering *S. flexneri* 2a O-antigen were obtained by density gradient centrifugation and observed by cryo-EM. OMVs, which are spherical with a bilayer membrane, could be observed. Flagellin was not detected due to the deletion of FliC and FljB proteins (**Fig. 2A**). The deletion of OmpA, OmpC, and OmpD proteins, the major porins in *Salmonella*, had no apparent effect on the conformation of the OMVs (**Fig. 2A**).

**Figure 2.**
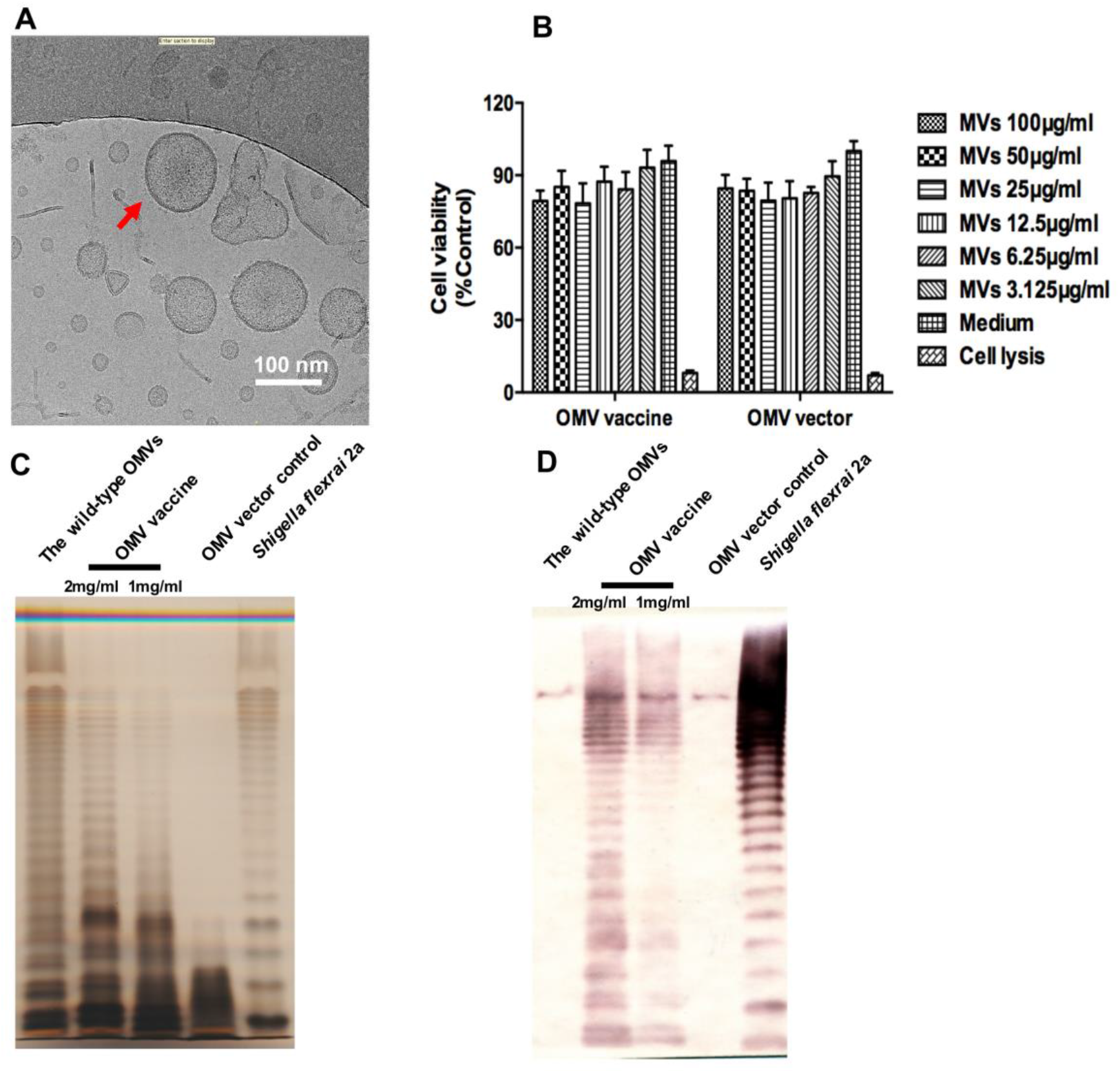
(A) OMVs derived from *S.* Typhimurium mutants delivering O-antigen were visualized by Cryo-EM. The image in B represent the white block in A enlarged 50 fold. The red arrow indicates a visible OMV. (B) Cytotoxic effect of OMVs derived from *S.* Typhimurium delivering *S. flexneri* 2a O-antigen polysaccharide and the parental mutant in RAW264.7 macrophages. Cells were incubated with the corresponding OMVs at the dose indicated. Cell viability was measured from the supernatant using a Multitox-Fluor Multiplex Cytotoxicity Assay. Supernatants from cells without OMV treatment and cell lysates acted as negative and positive controls, respectively. (C) Identification of OMVs derived from *S.* Typhimurium delivering *S. flexneri* 2a O-polysaccharide antigen by LPS profile and silver staining. (D) Identification of OMVs derived from *S.* Typhimurium delivering *S. flexneri* 2a O-polysaccharide antigen by Western blotting. The serum of *S. flexneri* 2a O-antigen was the first antibody to detect the band.

The safety of the OMV vaccine was also determined *in vitro* using a macrophage viability assay. RAW264.7 cells were treated with OMVs for 24 hours and their cytotoxicity examined using a MultiTox-Fluor Multiplex Cytotoxicity assay. The data demonstrated that neither the OMV vaccine or OMV vector elicited any apparent cytotoxicity. However, high concentrations of OMVs caused slight cell lysis (~75% cell viability) (**Fig. 2B**), indicating that, in spite of expressing the heterologous polysaccharide antigen, this OMV vaccine was sufficiently safe *in vitro* to subsequently evaluate it in *in vivo* experiments.

The biosynthesis of the *S. flexneri* 2a O-antigen in OMVs was determined by silver staining (**Fig. 2C**) and by Western blotting using anti-*S. flexneri* 2a serogroup serum (**Fig. 2D**). Deletion of the *rfbP* gene resulted in a change of LPS from smooth to rough phenotype in *S.* Typhimurium (**Fig. 2C**). The *S. flexneri* 2a O-antigen expressed by plasmid pQK018 were added and connected into the core polysaccharide using O-antigen ligase encoded by the *waaL* gene (**Figs. 2C and 1B**).

To further define the LPS bands from *S. flexneri* 2a O-antigen, Western blot analysis was performed. LPS samples from wild-type *S. flexneri* 2a and wild-type *S.* Typhimurium represented the positive and negative control, respectively (**Fig. 2D**). The LPS bands demonstrated a response to the anti-*Shigella* LPS serum, while other LPS from *Salmonella* displayed no bands on the nitrocellulose membranes (**Fig. 2D**). The results indicated that the O-antigen from *S. flexneri* 2a could be synthesized in OMVs derived by *S.* Typhimurium.

### Antibody response of *Salmonella*-derived OMVs delivering *Shigella* O-antigen

To evaluate the efficiency of the *Shigella* O-antigen-induced immune response due to their delivery in *Salmonella* OMVs, the mice were immunized with the recombinant OMV vaccine, *Salmonella* OMV vector or PBS control by intranasal or intraperitoneal administration. The concentration of antibodies, including IgG, S-IgA, and lung-wash IgA, against *S. flexneri* 2a LPS, was measured by quantitative ELISA. As shown in **Figure 3**, immunization with OMV vaccine resulted in a significantly higher anti-Shigella LPS IgG level than in the OMV vector group by both intranasal and intraperitoneal immunization. The intraperitoneal route of OMV vaccination elicited a greater serum antibody response compared with intranasal administration (*P* < 0.05) (**Figs. 3A and 3B**). Furthermore, mucosal immunity, as the first line of host defense, plays a critical role in the prevention of pathogenic infection. Secretory IgA (S-IgA) and lung-wash IgA are the most important factors in mucosal immunity against *Shigella* infection ^29^. Therefore, the concentration of S-IgA and lung-wash IgA in mice immunized were measured. It was found that the OMV vaccine delivering *S. flexneri* 2a O-polysaccharide antigen induced a significant mucosal antibody response against *Shigella* LPS (*P* < 0.05) (**Figs. 3C and 3D**). However, No mucosal antibody against LPS was detected in mice that were immunized intraperitoneally with OMVs after 8 weeks (data not shown).

**Figure 3.**
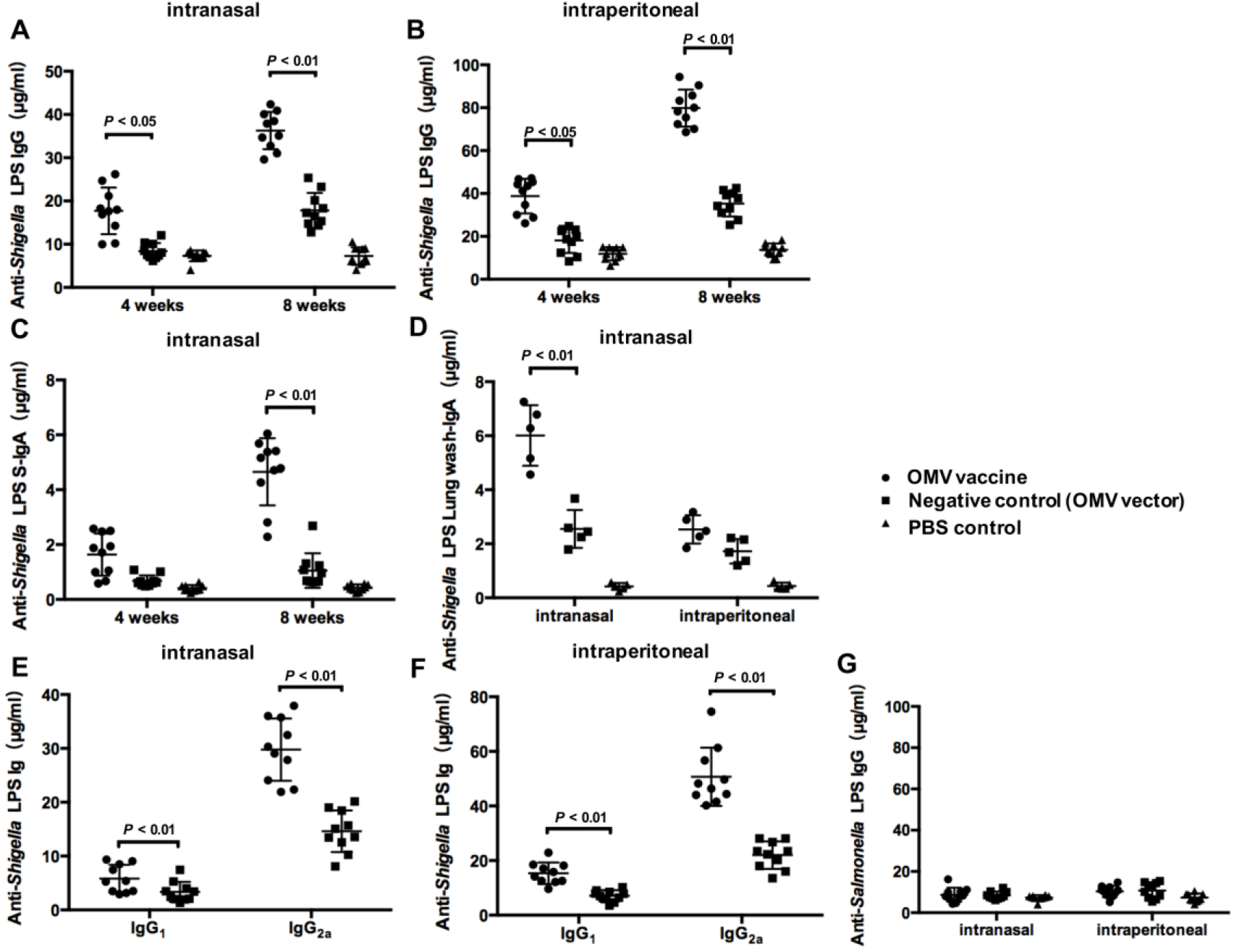
Humoral and mucosal antibody response in immunized and control mice after intranasal or intraperitoneal administration. *S. flexneri* 2a LPS was the coated immunogen. Serum IgG of mice immunized intranasally (A), serum IgG of mice immunized by intraperitoneal administration (B), total secretory IgA (S-IgA) of mice immunized intranasally (C), and lung wash-IgA from mice immunized intranasally (D) were quantified by ELISA. Each group represented 5 or 10 mice. The data represent exact concentrations of IgG or IgA antibodies quantified by a corresponding standard curve in individual sera from mice immunized intranasally or intraperitoneally using an OMV vaccine, OMV vector or PBS. Mice received a booster after 4 weeks and samples were collected 4 and 8 weeks after the first immunization. PBS-vaccinated mice represented the control group. Error bars represent variations among mice in each group.

### OMVs delivering *Shigella* O-antigen stimulated *Shigella* LPS-specific splenic lymophocyte proliferation

The priming of a cell-mediated immune response plays an important role in induction of protective immunity against *S. flexneri* infections. The proliferation of spleen cells in response to *Shigella* LPS and *S.* Typhimurium LPS stimulation was measured 7 days after primary immunization. The isolated spleen cells were cultured with *S. flexneri* 2a LPS or *S.* Typhimurium LPS, while the cells were compared with PHA culture. The results indicate that splenic cells extracted from mice with OMV vaccine administered intranasally and incubated with *S. flexneri* 2a LPS displayed significant lymphocyte proliferation similar to that of the PHA positive control group. In addition, no apparent proliferation of splenic lymphocytes incubated with S. Typhimurium LPS was observed (**Fig. 4A**). However, after extraction of spleen cells from mice immunized with OMV vaccine via intraperitoneal administration, cell proliferation was not observed whether incubated with *S. flexneri* 2a LPS or S. Typhimurium LPS (**Fig. 4B**).

**Figure 4.**
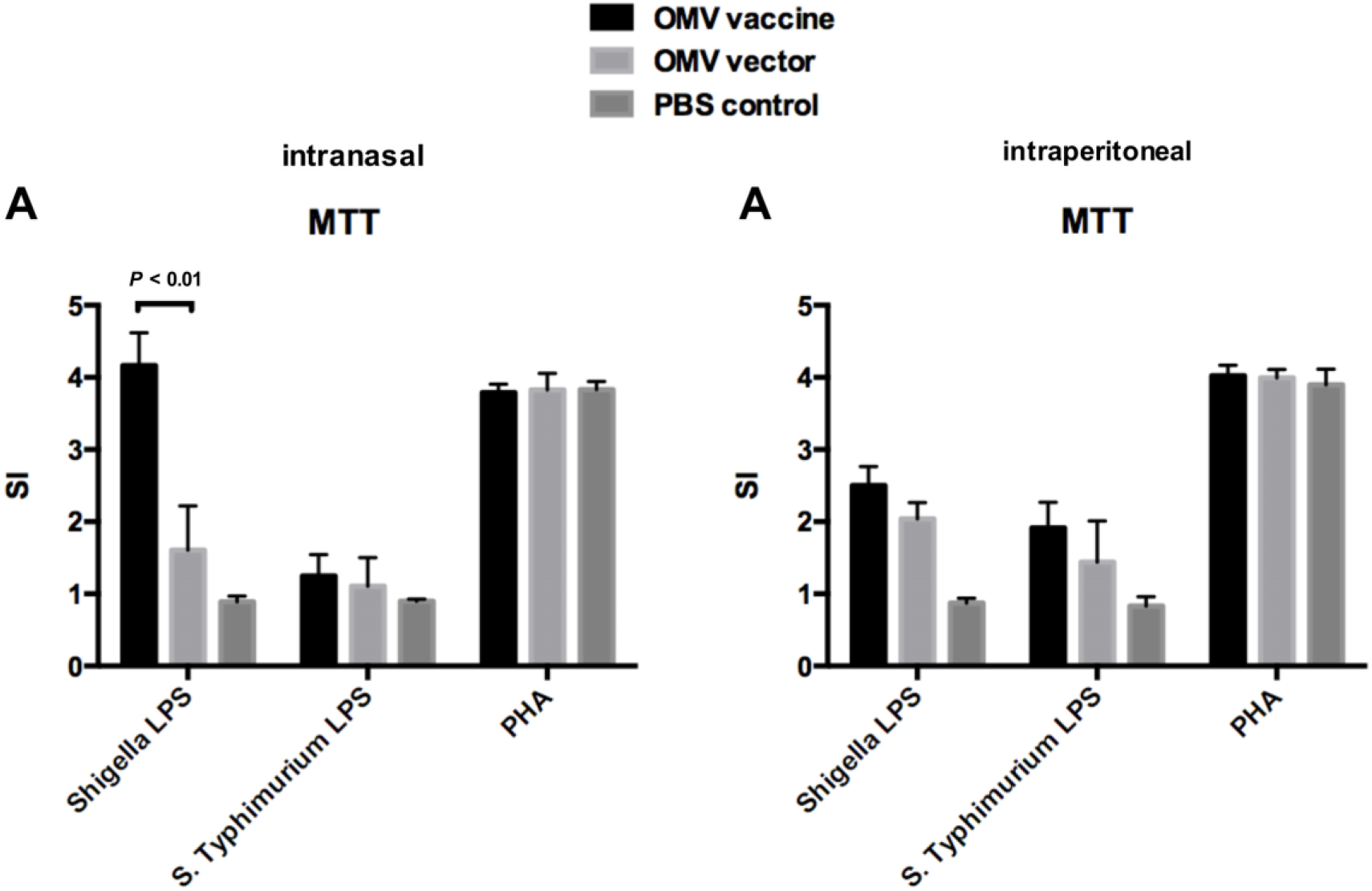
Splenic lymphocytes were isolated from mice which had been immunized for seven days by intranasal (A) or intraperitoneal administration (B). Each group consisted of 3 mice. Splenic lymphocytes were incubated with *S. flexneri* 2a LPS or S. Typhimurium LPS, while PHA was incubated as a positive control, to determine cell proliferation. The data represent proliferation of splenic lymphocytes in mice immunized intranasally or intraperitoneally with the OMV vaccine, OMV vector or PBS. Error bars represent variations between mice in each group. *P* < 0.01 compared with mice immunized with *Salmonella*-derived OMV vector (Negative control).

### Th1/Th2 polarization response to OMVs delivering *Shigella* O-antigen

Th1 and Th2 cells secrete a variety of cytokines that mediate an immune reaction. Therefore, the levels of IFN-γ, IL-12 (p40), IL-4, IL-13, IL-6, and TNF-α were measured to evaluate Th1/Th2 polarization. Splenocytes treated with the corresponding antigens were used to enhance immunity in order to detect cytokine levels. We found that IFN-γ, IL-12 (p40), IL-4, and IL-13 levels in spleen cells from both intranasal and intraperitoneal groups increased (**Figs. 5A-5D**). Of these, the levels of IFN-γ, IL-12 (p40), and IL-4 in cells treated with the OMV vaccine were higher than those in the OMV vector group (**Figs. 5A-5C**). As shown in **Fig. 5E**, IL-6 levels did not change significantly after a variety of treatments. Mouse splenocytes immunized with the intraperitoneal OMV vaccine displayed higher TNF-α levels than the intranasal route (**Fig. 5F**).

**Figure 5.**
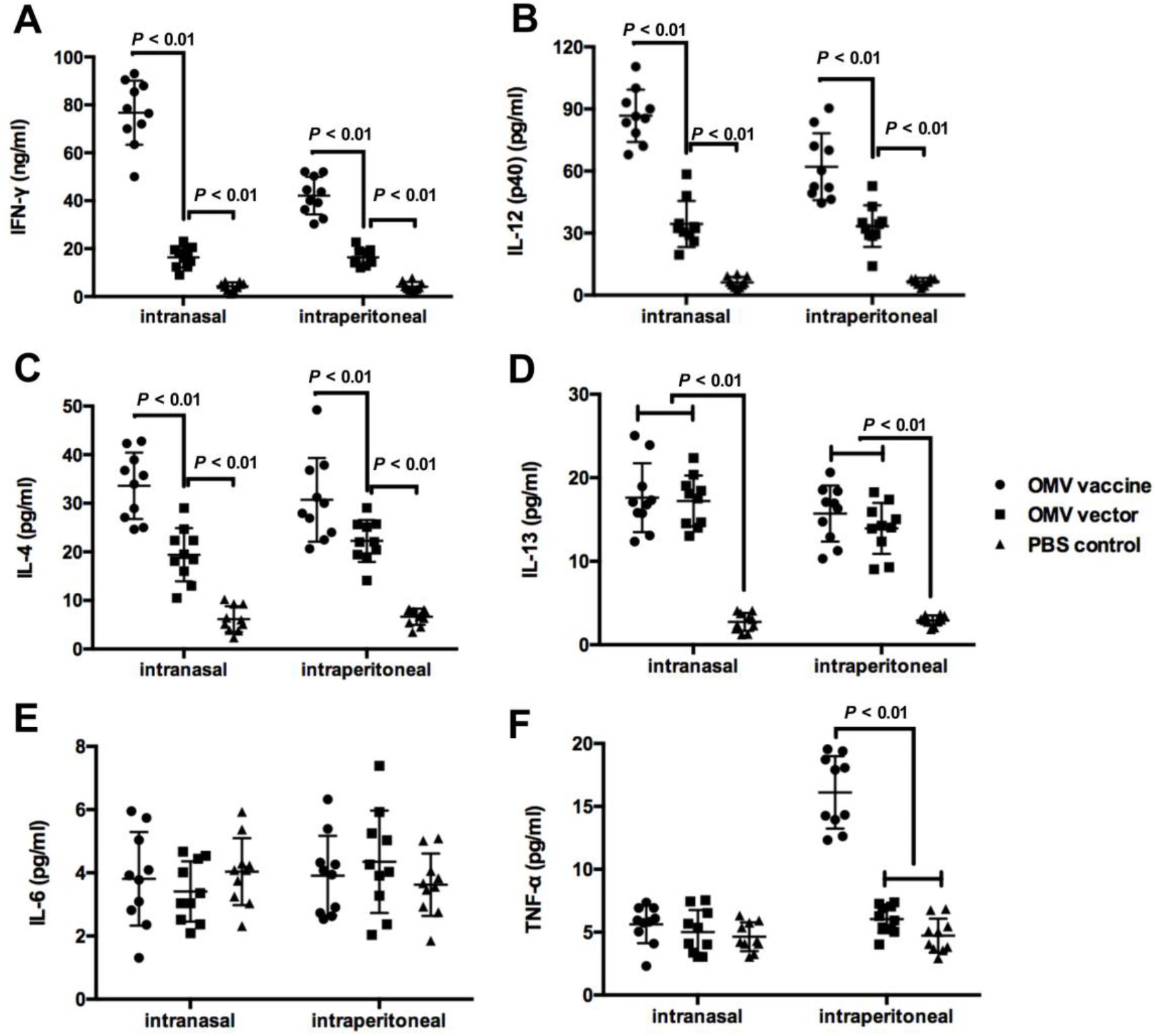
Splenocytes were isolated from 9 mice immunized intranasally or by intraperitoneal administration. Levels of cytokines were measured to evaluate the Th1/Th2 polarization effect. These data display the levels of cytokines in the splenocytes of mice immunized by intranasal or intraperitoneal administration with an OMV vaccine, OMV vector and PBS. After stimulation of the splenocytes with 6μg/ml LPS isolated from *S. flexneri* for 24 hours, levels of IFN-γ (A), IL-12 (p40) (B), IL-4 (C), IL-13 (D), IL-6 (E) and TNF-α (F) were measured by quantitative ELISA from the supernatants. Error bars represent variations between mice in each group. *P* < 0.01 compared with mice immunized with *Salmonella*-derived OMV vector (Negative control).

### SBA and opsonization assay

Aiming to evaluate specific bactericidal potential, sera from mice immunized with the recombinant OMV vaccine and *Salmonella* OMV vector were tested using an SBA. As presented in **Fig. 6A**, the intranasal OMV vaccine induced the secretion of antibodies with significantly higher SBA activity and >50% growth inhibition using a serum dilution of approximately 1:3,200 (*P* < 0.05). Although no statistical difference in SBA activity was found between the OMV vaccine and OMV vector immunized using intraperitoneal administration, a higher degree of inhibition was found in sera from mice intraperitoneally-immunized with the OMV vaccine compared those immunized with the OMV vector (**Fig. 6B**), indicating that *Salmonella*-derived OMVs delivering *Shigella* O-antigen polysaccharide elicited effective bactericidal activity.

**Figure 6.**
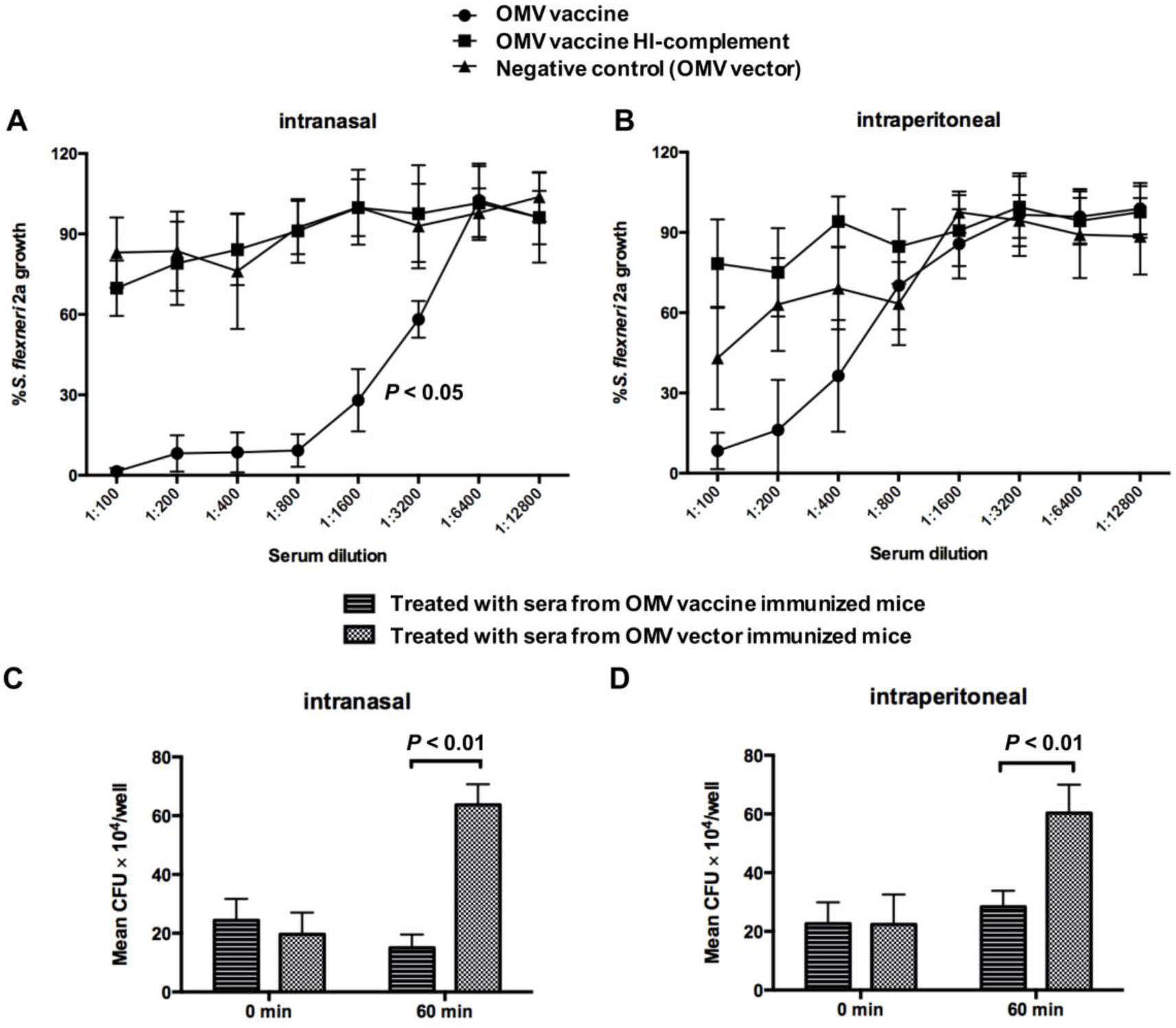
Bactericidal properties and Opsonization of sera from mice immunized with *Salmonella*-derived OMVs delivering *S. flexneri* 2a O-polysaccharide antigen *in vitro*. SBAs were performed using sera (8 weeks after first immunization) of mice in the OMV vaccine and OMV vector immunization groups, bacterial growth activity expressed in terms of serum dilutions. Mice immunized with OMVs by intranasal (A) or intraperitoneal (B) administration. The data represent the percentage CFU of *S. flexneri* 2a accounted for from each serum dilution with active/inactive complement compared with the CFU of the same serum dilutions with no complement. Error bars represent standard deviations. The opsonization assay was performed with mouse sera 8 weeks after the first immunization using the OMV vaccine and OMV vector. Mice immunized with OMVs by intranasal (C) or intraperitoneal (D) administration. Data represented the CFUs of *S. flexneri* 2a accounted for of each well with diverse-treated serum. Error bars represent standard deviations.

Furthermore, the ability of sera from mice immunized with the OMV vaccine to opsonize bacteria and enhance phagocytosis was determined using an isolated macrophage model. Bacterial uptake and death was observed in both intranasal and intraperitoneal groups (**Figs. 6C and D**), indicating that the sera of mice immunized with OMV vaccine displayed strongly-enhanced phagocytosis of *Shigella by* macrophages. *Salmonella*-derived OMVs that delivered *Shigella* O-antigen polysaccharide modulated the immune system of mice and participated in the clearing of pathogens (**Figs. 6C and D)**.

### Protection against *S. flexneri* 2a challenge

Mouse models have been typically used to evaluate specific protection against challenge with virulent *Shigella* infection ^19,20^. Five weeks after booster immunization, the mice were challenged with intranasal *S. flexneri* 2a at a lethal dose (approximately 10^6^ CFU, 100-fold LD_50_). This stringent challenge resulted in 100% mortality in PBS-immunized mice. However, mice immunized with intranasal OMV vaccine were 100% protected after the virulent *Shigella* challenge. In addition, the OMV vaccine given by intraperitoneal administration also provided significant protection to the mice (*P* < 0.05) (**Fig. 7A**). In particular, 1 or 2 mice immunized with OMV vector by both intranasal and intraperitoneal administration survived after the *Shigella* challenge, indicating that *Salmonella* OMV vector induced heightened innate immunity in these mice (**Fig. 7B**) ^45^.

**Figure 7.**
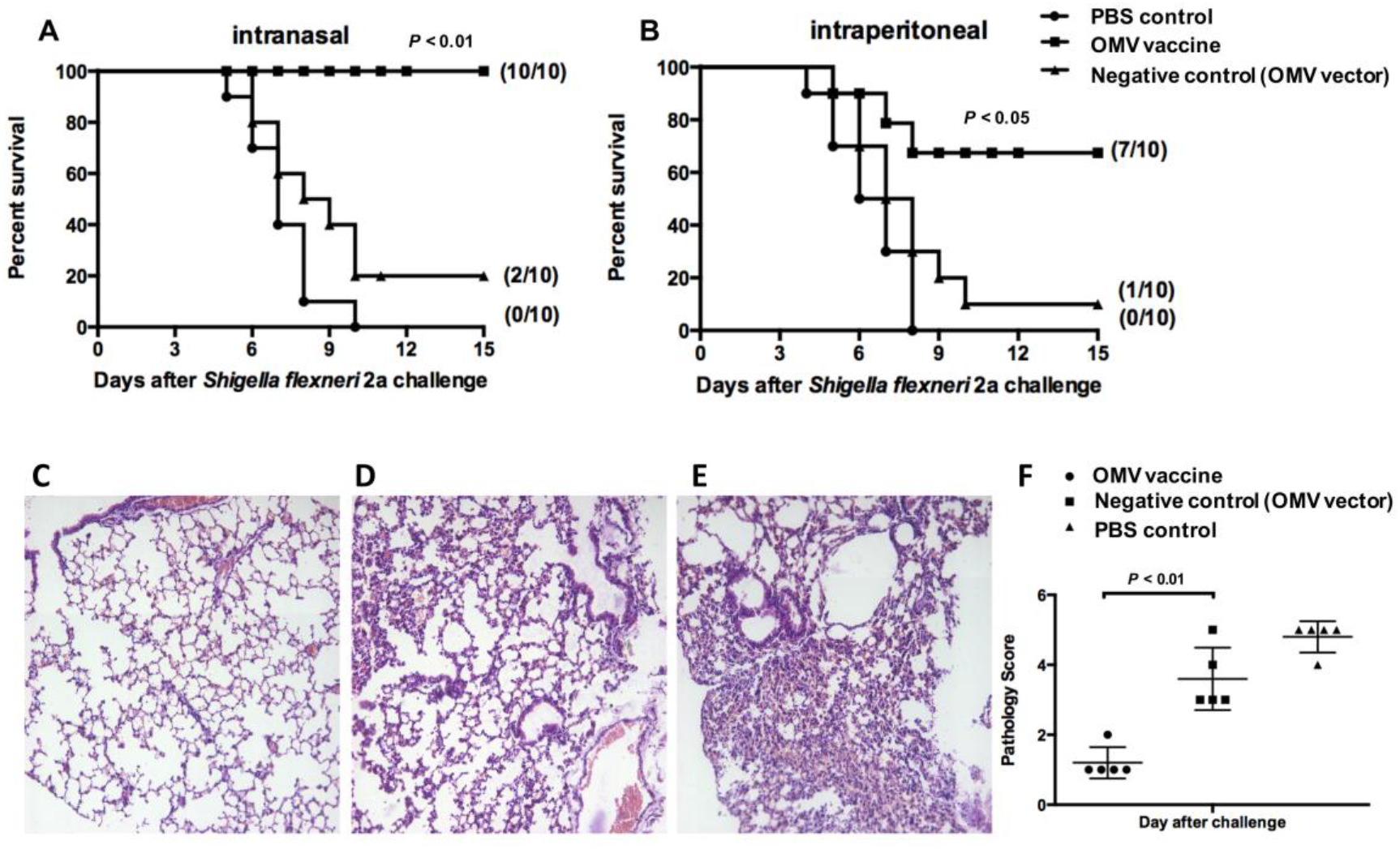
Intranasal (A) or intraperitoneal (B) immunization with *Salmonella*-derived OMVs delivering *S. flexneri* 2a O-polysaccharide antigen protected BALB/c mice against challenge with wild-type *S. flexneri* 2a by intranasal infection. Ten mice per group were immunized twice at 4-week intervals with the OMV indicated. Mice were challenged with 10^6^ CFU (~100-fold LD_50_) of *S. flexneri* 2a at 5 weeks, after booster immunization. Mortality was monitored for 3 weeks after challenge. The numbers in parentheses refer to the number of surviving mice and the total number of mice per group. *P* < 0.01 compared to mice immunized with *Salmonella*-derived OMV vector (Negative control). Hematoxylin and eosin-stained light micrographs (magnification x20) and pathology scores of histological sections from mice lungs. Lungs from intranasally immunized mice with OMVs delivering *S. flexneri* 2a O-polysaccharide antigen (C), 24 h after intranasal challenge with a lethal dose of *S. flexneri* 2a displaying normal airspaces, interstitium, and bronchioles. Lung from intranasally immunized mice with OMV vector (D) and from PBS-immunized mice (E) presenting severe pneumonia with vascular congestion, peribronchial and perivascular accumulation of inflammatory cells, including polymorphonuclear neutrophils and mononuclear cells. (F) Pathology scores of histological sections from mouse lungs. A score of 0 indicates no noticeable inflammation or lesions; a score of 1 indicates few or scattered foci affecting less than 10% of the tissue, typically with a few mild perivascular and/or peribronchial lymphoid aggregates; a score of 2 indicates frequent mild perivascular and/or peribronchial lymphoid aggregates, with or without occasional small foci of pneumonia, with overall inflammation affecting no more than 10 to 20% of the tissue; a score of 3 indicates moderate lesions, typically with abundant perivascular and peribronchial lymphoid infiltrates and multiple mild to moderate foci of pneumonia, with inflammation affecting approximately 20 to 30% of the tissue; a score of 4 indicates extensive pneumonia and marked inflammation affecting more than 30% of the tissue; and a score of 5 indicates extensive lesions with 50% of the tissue affected. *P* < 0.01 compared with mice immunized with *Salmonella*-derived OMV vector (Negative control).

To further confirm the protective efficacy of *Salmonella*-derived OMV vaccine, histological and pathological scores of mouse lung tissue after *S. flexneri* 2a challenge were determined. Lungs from mice immunized with the OMV vaccine displayed normal lung morphology (**Fig. 7C**). In comparison, mice immunized with the OMV vector displayed lung architecture with mild infiltration of neutrophils and mononuclear cells (**Fig. 7D**). However, mice immunized with PBS demonstrated significant perivascular and peribronchial inflammation, clearly characteristic of severe pneumonia (**Fig. 7E**). Statistical analysis of the pathological scores demonstrated consistency with histological analysis (**Fig. 7F**), indicating that the *Salmonella*-derived OMV vaccine may provide significant protection against virulent *S. flexneri* 2a infection in the mouse model of lung pneumonia.

## Discussion

Shigellosis is an important cause of morbidity and mortality among both children and adults, with *S. flexneri* the most frequently isolated species worldwide, and accounting for the majority of cases in developing countries ^46,47^. Vaccination appears to be a rational prophylactic approach in its control ^47^. In a recent study, many vaccine design strategies focused on serotype-specific *Shigella* LPS-associated O-antigen due to their ability to cross-protect against diverse serotypes, and O-polysaccharide antigen inducing a strong antibody responses against *Shigella* compared with other conserved invasion plasmid antigens (Ipas) ^2^. However, purified O-antigen is poorly immunogenic. Therefore, O-antigen is commonly coupled with protein carriers or functionally assembled on bacterial glycosyl carrier lipids to induce a stronger and longer-lasting immune response ^48,49^. An innovation of the present study was to combine the biosynthetic system of *Salmonella* glycoconjugates and OMVs, representing an efficient vaccine and delivery system ^31^, ideal as a vaccination against *Shigella*. For the first time, we report *S. flexneri* 2a O-polysaccharide antigen enzymatically conjugated to residues of the core-lipid structure using the oligosaccharyltransferase WaaL (RfaL) from *S.* Typhimurium (**Fig. 1B**). Glycoconjugates of the O-antigen were contained in purified *Salmonella* OMVs, as confirmed by silver staining and Western blot analysis (**Figs. 2C and 2D**).

To assemble and express the gene cluster for *S. flexneri* 2a O-polysaccharide antigen in *Salmonella*, the single-copy plasmid pYA3337 was used. A Gibson assembly kit was utilized to express and assemble the O-polysaccharide antigen, suitable for large DNA assemblies with heterologous antigen expression ^50^. Furthermore, to avoid foreign O-antigen becoming recognized by the host immune system via the carrier LPS ^51^, the *rfbP* mutation was introduced to truncate the carrier LPS. That mutation did not affect *S. flexneri* 2a O-antigen co-conjugating with the lipid A core (**Figs. 2C and 2D)**. In addition, a number of non-essential proteins, which could have affected the immune response of the polysaccharide antigen to the host immune system, were also omitted, such as FliC, FljB, OmpA, OmpC, and OmpD. Removal of flagellin has been shown to enhance the ability of OMVs to be recognized by host immune cells ^52^. In addition, our previous observations of cross-reactivity with the *Salmonella* OMV vaccine and vector demonstrate that the deletion of the major porin OmpACD enhanced cross-protection against *S.* Choleraesuis and *S.* Enteritidis infection (unpublished data), indicating that the *Salmonella* OMV vaccine, constructed for delivering the heterologous polysaccharide antigen, not only acted as a delivery antigen vaccine for the control of *Shigella* infection but also prevented infection and provided treatment for multiple serotypes of *Salmonella*. That might be because the major porin OmpACD elicited a strong but non-essential immune response that enhanced protective efficacy and delivered the heterologous antigen ^53,54^.

Safety is a primary consideration for vaccine design, and recent strategies have focussed on targeting purified or recombinant subunit vaccines because so far they represent the safest assured route for the development of a vaccine. However, safety is usually accompanied by low inherent immunogenicity. Therefore, OMV may represent an ideal approach that will balance safety with immunogenicity. The proposed approach involves OMVs naturally released from *S.* Typhimurium, and the present study established that these naturally purified OMVs have no cytotoxicity except at very high concentrations (**Fig. 3B**). However, such a high dose of OMV is not possible when vaccinating. Furthermore, it has been established that bacterial OMVs can enhance the immunogenicity of antigens that are naturally poorly immunogenic without the addition of adjuvants ^31,55^. Taken together, it is clear that the use of OMVs as a delivery carrier is of incomparable benefit.

In the present study, we demonstrated that both intranasal and intraperitoneal administration of *Shigella* O-antigen vaccine based on OMVs resulted in a robust LPS-specific antibody response in both systemic and mucosal compartments (**Figs. 3A and 3B**). The results of quantitative ELISA and challenge experiments presented here indicate that the *Salmonella* OMV-based vaccine designed to protect against *Shigella* challenge would be effective by both intranasal and intraperitoneal administration. However, the benefits of intranasal immunization appear to be greater, due to being a safer vaccination that avoids intragastric and intestinal degradation compared with intraperitoneal immunization ^56^. Furthermore, intranasal immunization induced greater levels of specific IgA in vaginal secretions and murine lungs (**Figs. 3C and 3D**). The data suggest that intranasal administration of the OMV vaccine could result in higher survival rates after infection with virulent *S. flexneri* 2a compared with theose receiving it through the intraperitoneal route (**Fig. 7**). Therefore, the data also imply that intranasal immunization may be preferable for vaccination to protect against pathogens that invade the respiratory tract, especially pneumonia caused by *S. flexneri* 2a.

An effective glycoconjugate candidate vaccine against virulent pathogens should be highly immunogenic and able to elicit bactericidal antibodies, enhancing phagocytosis of pathogens by macrophages ^41,42^. The results indicate that the O-antigen polysaccharide delivered by *Salmonella* OMVs has good immunological characteristics, with bactericidal potential, and providing opsonization (**Figs. 3, 5, and 6**). The study also indicates that use of *Salmonella* OMVs as a vector is the ideal strategy for developing a glycoconjugate vaccine.

*Shigella* preferentially invades the colon, eliciting a severe inflammatory response in humans, but not mice, which are generally resistant to invasion into the intestinal epithelium ^57^. However, previous studies have described the pathological histological features of lung tissue following *Shigella* infection in mice ^58^, localized infection manifesting as inflammation of the mucosal epithelium closely resembling intestinal shigellosis ^59^. Therefore, in the present study, the pulmonary mouse model was used to evaluate the efficacy of protection against *Shigella* infection using a *Shigella* O-polysaccharide antigen vaccine based on *Salmonella* OMVs. It was established that intranasal challenge of a lethal *S. flexneri* 2a *Shigella* infection caused clear symptoms of severe pneumonia in mice immunized with PBS (**Fig. 9**), while OMV vaccine-immunized mice received significant protection using the same challenge model, indicating that the pulmonary model was suitable for evaluation of the *Shigella* vaccine. The ideal OMV vaccine co-conjugated with *S. flexneri* 2a O-antigen has potential as an excellent candidate vaccine.

Future studies will determine the quantity of *S. flexneri* 2a polysaccharide antigen in *Salmonell*a-derived OMVs by quantification of LPS based on the Kdo method to demonstrate the efficacy of the co-conjugate by *Salmonella* biosynthesis. These results demonstrate the advantage of this strategy compared with traditional proteosome-LPS vaccines. While further studies will be required to establish whether an immune response against *Shigella* LPS could be enhanced by increasing the dose of OMV used in immunization and the number of doses that will be required of this OMV vaccine delivering O-polysaccharide antigen for sufficient immune protection against virulent *S. flexneri* 2a infection.

In summary, the study presented a novel strategy of delivering or co-conjugating a polysaccharide antigen using a biosynthetic *Salmonella* carrier and OMVs, which demonstrated immunogenicity and protective efficacy of this vaccine against virulent *S. flexneri* 2a infection in a murine pulmonary model that verified the potential of the vaccine using a *Shigella* O-antigen based on *Salmonella* OMVs. In addition, the results highlight the most suitable route of administration of this vaccine as intranasal for the control of Shigellosis. Finally, the study will assist in the development of polysaccharide co-cojugated vaccines and aid development of a multi-serotype vaccine platform using a vaccine design strategy based on OMV-based biosynthetic glycoconjugate.

## Acknowledgements

This study was supported by the National Natural Science Foundation of China (31760261), the Science and Technology Research Project of Jiangxi Provincial Education Department (60224), and Key research and development projects of Jiangxi Natural Science Foundation (20192BBG70067).

## Author contributions

Qiong Liu conceived and designed the experiments; Huizhen Tian, Biaoxian Li, Yuxuan Chen, Kaiwen Jie, Tian Xu and Zifan Song performed the experiments. Huizhen Tian, Qiong Liu and Xiaotian Huang analyzed the data; Huizhen Tian and Qiong Liu wrote the manuscript.

## Supplemental Figure

**Supplemental Figure 1.**
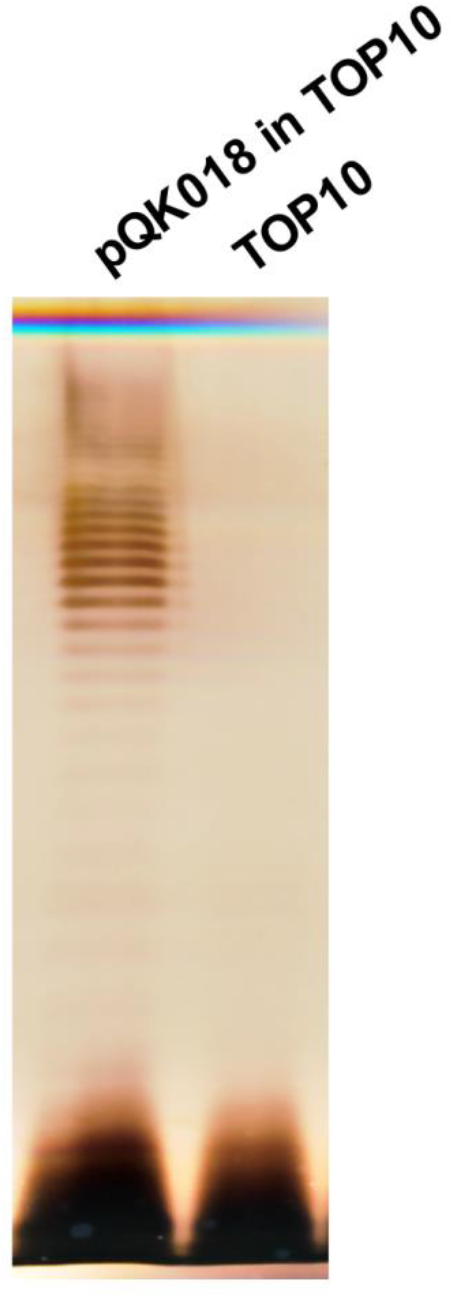
Identification of *E. coli* TOP10 delivering *S. flexneri* 2a O-polysaccharide antigen by LPS profile and silver staining.

